# CD16-158-Valine Chimeric Receptor T Cells Overcome the Resistance of KRAS-mutated Colorectal Carcinoma Cells to Cetuximab

**DOI:** 10.1101/588525

**Authors:** Roberto Arriga, Sara Caratelli, Giulia Lanzilli, Alessio Ottaviani, Carlo Cenciarelli, Tommaso Sconocchia, Giulio Cesare Spagnoli, Giandomenica Iezzi, Mario Roselli, Davide Lauro, Gianpietro Dotti, Soldano Ferrone, Giuseppe Sconocchia

**Affiliations:** Department of Systems Medicine, Endocrinology and Medical Oncology, University of Rome “Tor Vergata”, Rome, Italy; Institute of Translational Pharmacology, CNR, Rome, Italy; Otto Loewi Research Center, Chair of Immunology and Pathophysiology Medical University of Graz, Graz, Austria; Department of Surgery, Ente Ospedaliero Cantonale, Università della Svizzera Italiana, Lugano, Switzerland; Lineberger Comprehensive Cancer Center, University of North Carolina, Chapel Hill, NC, USA; Department of Surgery, Massachusetts General Hospital, Harvard Medical School, Boston, MA, USA

**Keywords:** Fc gamma CR T cells, polymorphisms, Immunotherapy, KRAS-mutated CRC, anti-EGFR mAb

## Abstract

KRAS mutation hinders the therapeutic efficacy of epidermal-growth-factor-receptor (EGFR) mAb (cetuximab and panitumumab)-based immunotherapy of EGFR+ cancers. Although, cetuximab controls KRAS-mutated cancer cell growth *in vitro* utilizing a NK cell-mediated antibody-dependent-cellular-cytotoxicity-(ADCC) mechanism, KRAS-mutated colorectal carcinoma (CRC) cells can still escape NK cell immunosurveillance. To overcome this limitation, we used cetuximab and panitumumab to redirect Fcγ chimeric receptor (CR) T cells against KRAS-mutated HCT116 CRC cells. We compared 4 polymorphic Fcγ-CR constructs including CD16^158F^-CR, CD16^158V^-CR, CD32^131H^-CR, and CD32^131R^-CR which were transduced into T cells utilizing retroviral transduction. Percentages of transduced T cells expressing CD32^131H^-CR (83.5±9.5) and CD32^131R^–CR (77.7.±13.2) were significantly higher than those expressing with CD16^158F^-CR (30.3±10.2) and CD16^158V^-CR (51.7±13.7) (p<0.003). CD32^131R^-CR T cells specifically bound soluble cetuximab and panitumumab. However, only CD16^158V^-CR T cells released significantly higher levels of interferon gamma (IFNγ=1145.5 pg/ml ±16.5 pg/ml, p<0.001) and tumor necrosis factor alpha (TNFα=614 pg/ml ± 21 pg/ml, p<0.001) than non-transduced T cells when incubated with KRAS-mutated HCT116 cells opsonized with cetuximab. Only CD16^158V^-CR T cells combined with cetuximab controlled the growth of HCT116 cells subcutaneously engrafted in CB17-SCID mice. These results suggest that CD16^158V^-CR T cells combined with cetuximab represent useful reagents to develop an effective immunotherapy of EGFR+KRAS-mutated cancer.

## INTRODUCTION

EGFR is overexpressed in several solid tumors ^1^. Upon binding to epidermal growth factor (EGF), EGFR triggers a series of signaling pathways supporting invasion and metastasis, and metastasis ^1^. The important role of EGFR in promoting cancer progression has provided the rationale to develop EGFR targeted mAb-based treatments ^2^.

EGFR-specific cetuximab is a chimeric IgG1 mAb preventing EGFR dimerization by stimulating its internalization and degradation. EGFR-specific panitumumab is a human IgG2 mAb interfering with EGF binding to its receptor. Cancer cells incubated with cetuximab or panitumumab undergo cell cycle arrest and apoptosis ^2^. However, a variety of EGFR+ cancer cells, including CRC cells, are insensitive to EGFR-specific mAbs since they carry RAS gene mutation(s) downstream of EGFR. Because of these mutations, cancer cells can bypass antitumor activities of both cetuximab and panitumumab. Lack of sensitivity of KRAS-mutated CRC cells to EGFR-specific mAb has serious clinical consequences since both cetuximab and panitumumab either have no effect on tumor growth or worsen CRC clinical course ^3,4^.

Increasing evidence suggests that this limitation can be overcome, at least for cetuximab, by taking advantage of its ability to mediate ADCC since EGFR+ cells, opsonized with cetuximab, undergo ADCC by activating the CD16 receptor expressed on NK cells ^5^. Nevertheless, cancer progression in patients is not arrested.

Failure to control cancer growth may be due to activation of evasion mechanism(s) from immune cells. However, in patients with cancer, NK cells, the major ADCC effector cells, show distinct functional defects and low ability to infiltrate solid tumors ^6–8^.

To restore the sensitivity of KRAS-mutated cancer cells to EGFR-specific mAbs, we investigated different strategies based on generation of extracellular CD16-CR linked to intracellular signaling and activating molecules ^9–12^. Because of NK cell limitations ^13^, we decided to use T cells, as effectors, since they easily infiltrate the tumor microenvironment and effectively protect hosts from cancer progression ^14^. Selected CD16-CR constructs have been transduced into T cells to redirect them by EGFR-specific mAbs toward EGFR+ cancer cells.

Other than NK cells, myeloid cells also mediate effector functions including proinflammatory cytokine production ^15,16^ and cytotoxic activity, including ADCC ^15,17^. Unlike NK cells, myeloid cells have the exclusive property to recognize Fc fragments of IgG2 antibodies complexed with the corresponding antigens on target cells, utilizing the Fc receptor CD32 and triggering ADCC activation ^18^. Both CD16 and CD32 are polymorphic and their polymorphisms influence their binding to IgG Fc fragments ^19^.

Still unknown is whether CD32 and CD16 polymorphisms impact the anti-tumor activity of Fcγ-CR T cells against KRAS-mutated CRC cells. The goal of this study is to compare the ability of polymorphic CD16-CR and CD32-CR to inhibit KRAS-mutated CRC cell proliferation and tumor progression *in vitro* and *in vivo*.

## MATERIALS AND METHODS

### Antibodies, reagents and cell lines

Fluorescein isothiocyanate (FITC)-conjugated mouse anti-human CD3 (cat.555332), phycoerythrin (PE)-conjugated mouse anti-CD16 (cat.555407), PE-conjugated mouse antihuman CD32 (cat. 550586), mouse anti-human CD3 (cat. 555329), CD28 (cat.555725), CD32 (8.26) (cat. 557333), and CD16 (3g8) (cat. 556617) were purchased from BD Bioscience (San Jose, CA, USA). Cetuximab (Erbitux 5mg/ml) and panitumumab (Vectibix 20mg/ml) were purchased from Merck Serono (Darmstadt, Germany) and from Amgen (Thousand Oaks, CA, USA), respectively. 3-(4,5-Dimethylthiazol-2-Yl)-2,5-Diphenyltetrazolium Bromide (MTT) was purchased from Sigma-Aldrich (Saint Louis, MO, USA) and GeneJuice^®^ Transfection Reagent (Novagen) from Millipore (Burlington, MA, USA). Human recombinant interleukin-7 (IL-7) and interleukin-15 (IL-15) were purchased from Peprotech (London, UK) and Lipofectamine 2000 from Life Technologies (Carlsbad, CA, USA). Retronectin (Recombinant Human Fibronectin) was purchased from Takara Bio (Saint-Germain-en-Laye, France). Dulbecco’s Modified Eagle’s Medium (DMEM), Iscove’s Modified Dulbecco’s Medium (IMDM), RPMI 1640 medium, FBS, L-glutamine and penicillin/streptomycin were purchased from Thermo Fisher Scientific (Waltham, MA, USA). Complete media (CM) were supplemented with 10% FBS, 2 mM L-glutamine, 0.1 mg/mL streptomycin, and 100U/ml penicillin. Mycoplasma-free, KRAS-mutated HCT116 cells ^20^, provided by Giulio Spagnoli (University of Basel, Basel, CH) were maintained in RPMI 1640, CM. Cells were authenticated on November 21th, 2018 by PCR-single-locus-technology (Eurofins, Ebersberg, Germany). Cells were kept in culture for a maximum of 4 to 8 passages.

### Fc_γ_ chimeric receptors

Generation of CD16^158F^-CD8α-CD28-CD3ζ CR has previously been described ^12^. Extracellular region of CD32A^131R^ was amplified by reverse-transcriptase polymerase chain reaction (RT-PCR) from RNA extracted from peripheral blood mononuclear cells (PBMCs) utilizing the following primers: forward 5’-GAGAATTCACCATGACTATGGAGACCCAAATG-3’ and reverse 5’-CGTACGCCCCATTGGTGAAGAGCTGCC-3’ (Thermo Fisher Scientific, Waltham, MA, USA). PCR product was fused by restriction enzyme-compatible ends with the CD8α-CD28-CD3ζ domain contained in the pcDNA3.1/V5-His TOPO TA (Invitrogen, Carlsbad, CA, USA). CD16^158F^-CR and CD32^131R^-CR were subcloned into NcoI and MluI sites of the SFG retroviral vector. CD16^158V^ and CD32A^131H^ were assembled by using synthetic oligonucleotides. Fragments were separately inserted into SFG vector and sequenced by GeneArt Gene Synthesis team (Invitrogen-Thermo Fisher Scientific, Regensburg, Germany).

### Retrovirus production and T cell transduction

Retroviral supernatants were obtained by transient transfection of 293T cells, with Peg-Pam plasmid encoding the Moloney murine leukemia virus gag and pol genes, and RDF plasmid encoding the RD114 envelope and the CD32^131R^-, CD32^131H^-, CD16^158F^- or CD16^158V^-CR SFG retroviral vectors, using GeneJuice reagent. Forty-eight and 72h posttransfection, retrovirus-containing supernatants were harvested, filtered, snap frozen, and stored at −80°C until use. To generate Fcγ-CR T cells, PBMCs (0.5×10^6^ PBMCs/ml) were cultured for 3 days in non-tissue culture treated 24-well plates pre-coated with 1 μg/ml anti-CD3 and 1 μg/ml anti-CD28 mAbs in the presence of 10 ng/ml IL-7 and 5 ng/ml IL-15. Viral supernatants were placed on retronectin-coated non-tissue culture treated 24 well plates and spun for 1.5h at 2000xg. Activated T cells were seeded into retrovirus loaded-plates, spun for 10’, and incubated for 72h at 37°C in 5% CO_2_ atmosphere. After transduction, T cells were expanded in RPMI 1640 CM supplemented with 10 ng/ml IL-7 and 5 ng/ml IL-15.

### *In vitro* tumor cell viability assays

Antitumor activity of Fcγ-CR T cells *in vitro* was evaluated by MTT assays. Tumor target cells (7×10^3^/well) were seeded in 96-well plates and Fcγ-CR T cells (35×10^3^/well) were added in the presence or absence of (3 μg/ml) cetuximab or panitumumab. Following a 48-72h incubation at 37°C, non-adherent T cells were removed and 100 μl/well of fresh medium supplemented with 20 μl of MTT (5 mg/ml) were added to adherent cells and incubation was continued for 3h at 37°C. Following a 3h incubation at 37°C, supernatants were removed and 100 μl of dimethyl sulfoxide (DMSO) were added to each well. Absorbance was measured at 570 nm.

### Cytokine release

Two-fold dilutions of a Fcγ-CR T cell suspension (1×10^5^/100μl/well) were added to 96 well plates in triplicates. Then, EGFR+, KRAS-mutated HCT116 cells (2×10^4^/100μl/well) were added at a 5:1 E: T ratio in the presence or absence of 3μg/ml cetuximab or panitumumab. Supernatants were collected following a 48h incubation at 37°C and IFNγ and TNFα levels were measured by ELISA (Thermo Fisher Scientific, Waltham, MA, USA).

### Flow cytometry

Fcγ-CR expression levels on engineered T cells were assessed by flow-cytometry upon incubation for 30min at 4°C with FITC-conjugated mouse anti-human CD3, PE-conjugated mouse anti-human CD32 or PE-conjugated mouse anti-human CD16 mAbs. Cells were analyzed by 2-laser BD FACSCalibur (Becton Dickinson, Franklin Lakes, NJ, USA) flow cytometer. Results were evaluated utilizing Tree Star Inc. FlowJo software.

### Xenograft mouse model

*In vivo* experiments were performed in accordance with the directive 2010/63/EU and authorized by the Italian Ministry of Health (186/2016-PR). Antitumor activity of CD16^158V^-CR T cells with or without cetuximab was assessed using 8-week-old male CB17-SCID mice (CB17/lcr-PrkdcSCID/lcrlcoCrl, Charles River Laboratories, Lecco, Italy, Cat.CRL:236, RRID: IMSR CRL:236), 12-18gr body weight, engrafted with KRAS-mutated HCT116 CRC cells. Mice were housed in temperature-controlled rooms with 12h light/dark cycle and free access to sterile water and autoclaved standard chow diet (4RF25; Mucedola, Milan, Italy). Endogenous NK cell activity was suppressed by intraperitoneal injection of 20μl rabbit anti-asialo-GM1 antibody (Wako, Chemicals, Richmond, VA, Cat.986-10001). Mice received anti-asialo-GM1 antibody on days −3, 0, +14, and +21 since tumor cell engraftment. On day 0, mice were grafted subcutaneously in the right flank, with 1×10^6^ HCT116 cells and then randomly separated into 4 groups (5 mice per group). Group 1 received only HCT116; group 2 cetuximab (150μg); group 3 HCT116 and CD16^158V^-CR T cells and group 4 HCT116, CD16^158V^-CR T cells and cetuximab (150 μg). Effector cells were administered at a 5:1 E:T ratio. Tumor volumes were measured every 3 days with caliper and calculated using the formula: TV (cm^3^) = 4/3p r^3^, where r= (length + width)/4. When tumor volume reached 2 cm^3^ mice were sacrificed.

### Statistical analysis

Results were analyzed by paired-*T*-test, Mann-Whitney test and two-way analysis of variance (ANOVA) followed by Bonferroni’s multiple-comparison, as necessary. Disease-free survival (DFS) was evaluated by log-rank-(Mantel-Cox) test. Differences were considered significant with *p*-values < 0.05.

## RESULTS AND DISCUSSION

To enhance anti-tumor potential of EGFR-specific mAbs, we generated CD16 and CD32-CR (fig.1A). We successfully expressed all indicated Fcγ-CR into transduced T cells (fig.1B). However, T cell transduction of CD32^131H^-CR (83.5%±9.5%) and CD32^131R^-CR (77.7%±13.2%) was significantly more effective than that of CD16^158V^-CR (51.7%±13.7%) and CD16^158F^-CR (30.3%±10.2%) (p<0.003 fig.1C). Taking in consideration that all the retroviruses are packaged using the same methodology and that the titer of the produced viruses are similar, these results may suggest that in human T lymphocytes polymorphic CD32-CR are more stably expressed as compared to polymorphic CD16-CR even if the underlying mechanism(s) remain to be explored. However, Cheeseman et al., provided evidence that hematopoietic cells tend to express CD32 more efficiently than CD16 on their surfaces ^21^.

**Figure 1.**
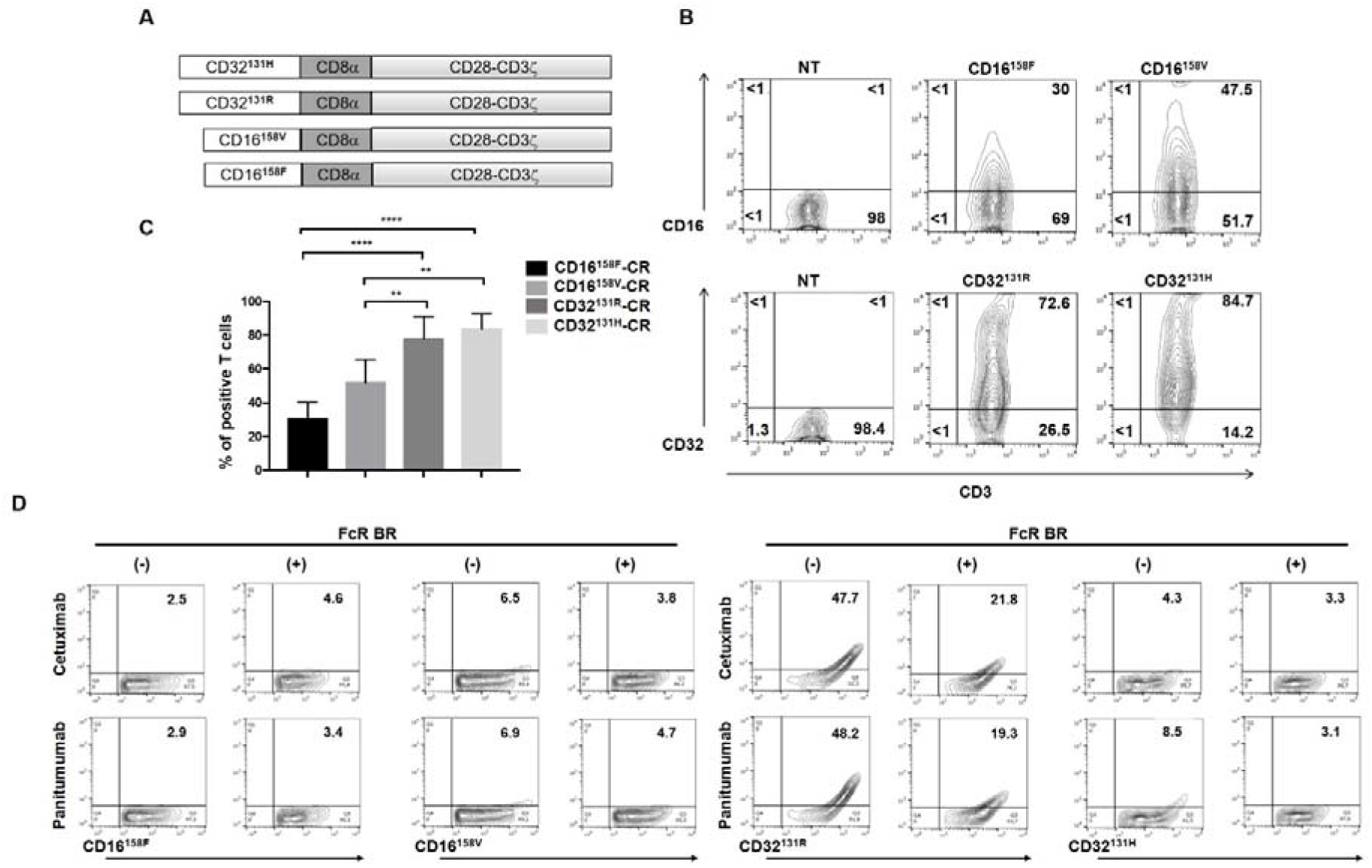
Molecular structure and functional features of Fcγ-CRs. Panel A shows a schematic representation of the gene constructs encoding the indicated Fcγ-CRs. The extracellular domain of high and low-affinity CD32-CRs and CD16-CRs incorporate the transmembrane CD8α and CD28/CD3ζ chain. Panel B reports a representative experiment showing the expression of Fcγ-CRs on the surface of T cells 48h after retroviral transduction. Percentages of cells expressing the transgenes are shown in the quadrants. Panel C shows cumulative percentages of positive T lymphocytes for each Fcγ-CR 48h following viral infection of cells from five healthy donors. Asterisks: ****= p< 0.0001, **= p<0.003. Panel D shows the binding of soluble cetuximab and panitumumab on the surface of Fcγ-CR T cells. T lymphocytes expressing each CD16-CR and CD32-CR polymorphisms were incubated with 3 μg/ml of either cetuximab or panitumumab for 30min at 4°C with or without FcR BR. Cells were then washed and stained with a FITC-conjugated mouse anti-human IgG antibody. Binding of cetuximab and panitumumab was then evaluated by flow cytometry. FcR BR: Fc receptor blocking reagent.

To evaluate Fcγ-CR T cell antibody binding capacity, polymorphic CD32-CR and CD16-CR T cells were incubated with cetuximab or panitumumab, for 30min, at 4°C. CD32^131R^-CR T cells efficiently bound both cetuximab and panitumumab while CD32^131H^-CR T cells only displayed a minimal binding for panitumumab. In contrast, CD16^158V^-CR T cells and CD16^158F^-CR T cells failed to bind both cetuximab and panitumumab (fig.1D). Binding of cetuximab and panitumumab to CD32-CR T cells was highly specific and promptly inhibited in the presence of Fc receptor blocking reagent (FcR BR) (fig.1D). CD16 and CD32 binding affinity for IgG is known to be influenced by their polymorphisms. Presence of valine instead of phenylalanine at position 158 of CD16 (CD16^158V/V^) and presence of histidine instead of arginine at position 131 of CD32 (CD32^131H/H^) enhance IgG binding affinity of these receptors ^19,22^. CD16-CR has also been produced in other laboratories while, to the best of our knowledge, CD32-CR has not. CD16^158V^ has preferentially been utilized for *in vitro* and *in vivo* studies ^9–11,23^. However, Kudo et al. showed that CD16^158V^-CR has significantly higher affinity for rituximab than CD16^158F^-CR. Here, neither CD16^158V^-CR nor CD16^158F^-CR showed significant binding affinity for soluble cetuximab and panitumumab. Inability of our CD16-CR to bind cetuximab, might be related to structural differences of endodomains since we used CD28 while Kudo et al. used 4-1BB. CD32^131R^-CR bound soluble cetuximab and panitumumab with higher affinity than CD32^131H^-CR. As observed with CD16-CR, structural differences of CD32-CR endodomains may also influence their IgG binding affinity.

We next asked whether polymorphic Fcγ-CR T cells recognize KRAS-mutated HCT116 cells opsonized with EGFR-specific mAb and affect their viability. Transduced and non-transduced T cells were incubated, for 72h, at 37°C with KRAS-mutated HCT116 cells with or without cetuximab or panitumumab. IFNγ and TNFα production was measured in culture supernatants. Only CD16^158V^-CR T cells, combined with cetuximab but not panitumumab, produced levels of both IFNγ (1145.5±16.5 pg/ml) and TNFα (614±21 pg/ml) significantly higher than those released by the other Fcγ-CR T cells or nontransduced T cells (fig.2A).

**Figure 2.**
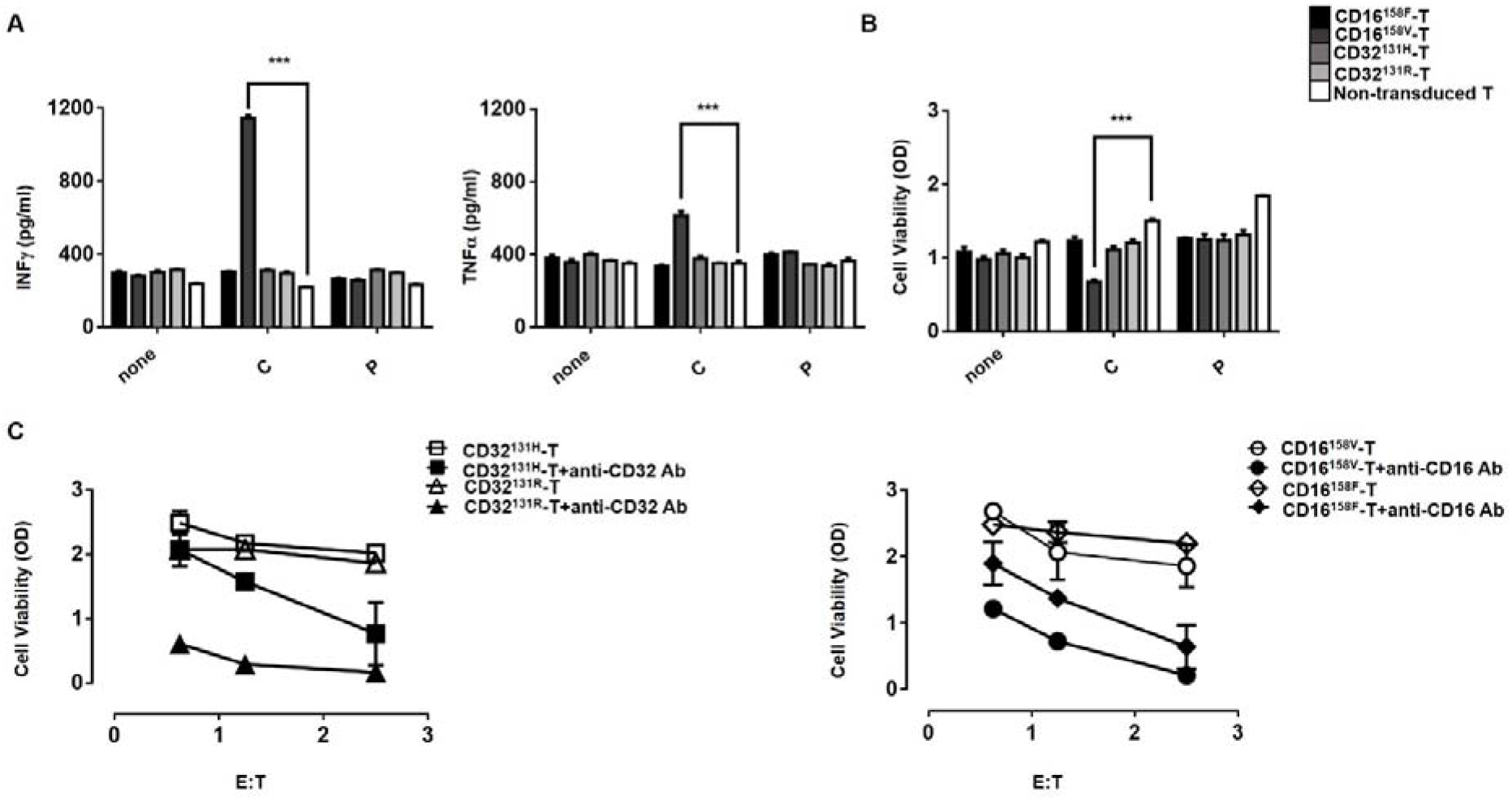
Recognition of HCT116 CRC cells by CD16^158V^-CR T cells in combination with cetuximab leads to proinflammatory cytokine production and HCT116 cell elimination. Panel A shows IFNγ and TNFα levels in supernatants of HCT116 cells incubated for 48h at 37 °C with the indicated Fcγ-CR T cells, in the presence or absence of cetuximab (3 μg/ml) or panitumumab (3 μg/ml) both at a 5:1 E: T ratio. Panel B shows viability, as evaluated by MTT assays of HCT116 cells incubated for 48h at 37°C with the indicated Fcγ-CR T cells with or without cetuximab or panitumumab at a 5:1 E: T cell ratio: C: cetuximab, P: panitumumab, white bars: non-transduced T cells. Asterisks indicate a p-value < 0.001. The figure reports cumulative data, with mean±SD values, of HCT116 cell viability obtained by using effector cells from 5 different donors in independent experiments. Panel C shows the results of redirected assays on the viability of stably transfected CD32+HCT116 cells, as measured by the MTT assay. Fcγ-CR T cells were incubated for 3 days, at 37°C with CD32+HCT116 in the presence or absence of anti-CD16 mAb (3mg/ml) (left panel) with or without anti-CD32 mAb (3 μg/ml) (right panel) at the indicated E: T cell ratio. Viability of HCT116 target cells was then measured as described in the method section. Asterisks indicate p<0.001.

Then, we tested whether polymorphic Fcγ-CR engineered T cells could impair viability of KRAS-mutated HCT116 cell opsonized with cetuximab or panitumumab, utilizing an ADCC mechanism. Figure 2B shows that KRAS-mutated HCT116 cell viability, expressed in optical density (OD), was significantly reduced following a 48h incubation at 37°C with CD16^158V^-CR T cells and cetuximab (0.67-OD±0.03-OD) as compared to nontransduced T cells (1.5-OD±0.03-OD). In contrast, no change in viability was detected when cetuximab was replaced with panitumumab. Both polymorphic CD32-CR T cells and CD16^158F^-CR T cells with or without anti-EGFR mAbs, failed to impair KRAS-mutated HCT116 cell viability. These data indicate that the efficient recognition of cetuximab-opsonized cancer cells by CD16^158V^-CR T cells leads to the activation of effector functions including production of proinflammatory cytokines and impairment of KRAS-mutated HCT116 cell viability. Therefore, CD16^158V^-CR T cells restore the ability of cetuximab to target KRAS-mutated HCT116 CRC cells by an immune-mediated mechanism *in vitro*.

Since these data challenged the effector potential of CD16^158F^-CR T cells and CD32-CR T cells, we tested them on KRAS-mutated HCT116 cells stably transfected with CD32 (Caratelli et al. unpublished data) in redirected ADCC assays. Although to different extents, all CR-engineered T cells, at different E: T ratio, significantly reduced the viability of FcγR+HCT116 cells in the presence of 3g8 (anti-CD16) or 8.26 (anti-CD32) agonistic mAb (Figure 2C). These data indicate that all engineered T cells are fully competent effector cells. Thus, although transgenic CD32 binds soluble mAbs and is able to provide a cytotoxic signal, CD32-CR T cells are unable to produce pro-inflammatory cytokines in coculture with opsonized KRAS-mutated HCT116, and to impair their viability. Therefore, soluble mAb binding and redirected killing do not represent effective surrogate assays for the ability of transduced T cells to elicit mAb-mediated effector functions targeting tumor cells. Notably, affinity of IgG1 cetuximab for CD32 is low and panitumumab mediates ADCC only in the presence of myeloid cells ^18^.

*In vivo* antitumor potential of CD16^158V^-CR T cells was then assessed. Transduced cell ability to impair KRAS-mutated HCT116 viability was tested prior to their administration to experimental animals. CD16^158V^-CR T cells were incubated at 37°C with target cells and cetuximab or panitumumab while non-transduced T cells were used as a negative control. Following 72h incubation, KRAS-mutated HCT116 viability was assessed by the MTT assay. Figure 3A confirms that co-culture with CD16^158V^-CR T cells, combined with cetuximab, significantly affected HCT116 cell viability. Instead, CD16^158V^-CR T alone or with panitumumab or non-transduced T cells were completely ineffective. We then engrafted CB17-SCID mice subcutaneously with KRAS-mutated HCT116 cells. One hour later CD16^158V^-CR T cells, with or without cetuximab, were injected in proximity of the injection site of HCT116 cells. Figure 3B shows tumor volumes of HCT116 on day 64 postinjection. CD16^158V^-CR T cells combined with cetuximab, significantly protected mice from tumor growth. DSF of treated animals is reported in figure 3C. Interestingly, tumor growth was also significantly delayed in the group of mice receiving CD16^158V^-CR. These results indicate that CD16^158V^-CR T cells combined with cetuximab and, to a lesser extent, even without cetuximab, exert significant anti-tumor activity *in vivo*.

**Figure 3.**
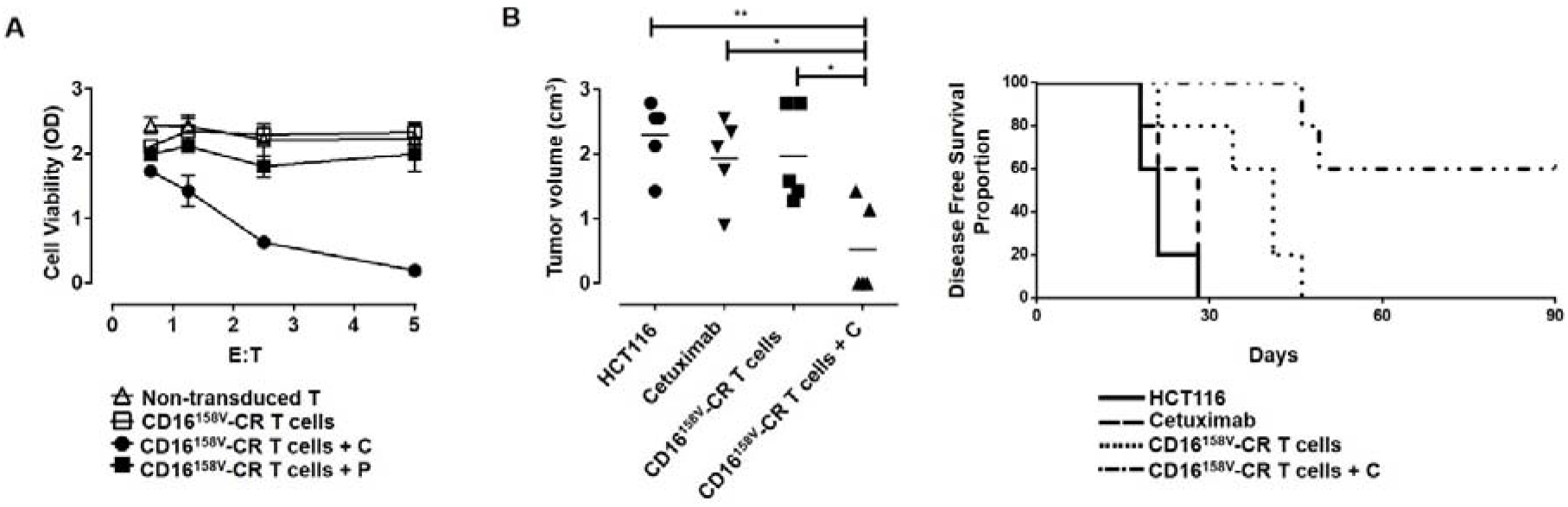
CD16^158V^-CR T cells in combination with cetuximab protect CB17 SCID mice from subcutaneous growth of KRAS-mutated HCT116 cancer cells. Panel A: HCT116 cells were incubated for 72h, at 37°C, in the presence or absence of CD16^158V^-CR T cells at the indicated E: T ratios with or without cetuximab (C) or panitumumab (P) both at the concentration of 3μg/ml. Non-transduced T cells were used as a control. HCT116 cell viability was evaluated by MTT assay. Values are expressed as mean± SD. Data are analyzed by a two-way ANOVA test. Panel B: four groups of CB17 SCID mice (N=5 per group) were injected with HCT116 cells (1×10^6^) subcutaneously, in the right flank. Following HCT116 injection, three groups of mice were injected, in the area adjacent to HCT116 injection, with 150 μg of cetuximab (group 2), 5×10^6^ CD16^158V^-CR T cells (group 3), and 5×10^6^ CD16^158V^-CR T cells plus 150 μg of cetuximab (group 4). HCT116 cell growth was then monitored. Left panel shows a scatter plot analysis of tumor volumes resulting from subcutaneous injection of KRAS-mutated HCT116 cells 64-days postinjection. Right panel shows DFS analysis, as evaluated by the Log-rank (Mantel-Cox) test. Asterisks: *= p-value 0.02, **= p-value 0.001, **** = p-value 0.0001.

This is the first study demonstrating the ability of CD16^158V^-CR combined with cetuximab can control KRAS-mutated cancer cell growth *in vitro* and *in vivo*. Our results by using the same KRAS-mutated HCT116 cells and at an equivalent reverse ADCC potential of Fcγ-CR utilized, the superior activity of CD16^158V^-CR may reflect its optimal interaction with the cetuximab Fc fragment ^5^. Taken together, these data contribute to a repositioning of currently available anti-EGFR therapeutic mAbs in the treatment of insensitive tumors, and pave the way toward innovative immunotherapies targeting KRAS mutated cancers.

## Abbreviations

EGF: epidermal growth factor
EGFR: epidermal-growth-factor-receptor
mAb: monoclonal antibody
CRC: colorectal carcinoma
ADCC: antibody-dependent-cellular-cytotoxicity
CR: chimeric receptor
INFγ: interferon gamma
TNFα: Tumor necrosis factor alpha
FITC: fluorescein isothiocyanate
IL-7: interleukin-7
IL-15: interleukin-15
RT-PCR: reverse-transcriptase polymerase chain reaction
PBMCs: peripheral blood mononuclear cells
DMEM: Dulbecco’s Modified Eagle’s Medium
IMDM;: Iscove’s Modified Dulbecco’s Medium
CM: complete media
DMSO: dimethyl sulfoxide
DSF: disease-free survival
FcR BR: Fc receptor blocking reagent
OD: optical density

## ACKNOWLEDGMENTS

This work was supported by the Italian Association for Cancer Research (AIRC) under grant IG17120 to GS and by NIH grants CA216114 and CA231766 to SF. We thank Marta Coccia and Antonio Rossi for technical assistance

